# Effect of Futsal-Based Game Training on Performance, Self-Efficacy, Motivation, and Exercise Addiction in Adolescent Non-Athlete Girls

**DOI:** 10.1101/2024.09.28.615615

**Authors:** Sahar Beik, Jalal Dehghanizadeh

**Affiliations:** Dep of Motor Behavior and Sport Management, Urmia University, Urmia, Iran

**Keywords:** futsal, game-based training, performance, self-efficacy, motivation, exercise addiction, adolescents

## Abstract

This study examined the effects of futsal-based game training on performance, self-efficacy, motivation, and exercise addiction in adolescent non-athlete girls. Thirty female students with no prior futsal experience were randomly assigned to either a traditional training group (n=15) or a game-based training group (n=15). Performance was assessed using a futsal skills test, while self-efficacy, sports motivation, and exercise addiction were measured via questionnaires. Both groups underwent separate 12-week training protocols. Analysis of covariance revealed significant differences between the game-based and traditional training groups in performance (P=0.0001), self-efficacy (P=0.0001), and sports motivation (P=0.0001). However, no significant difference was observed in exercise addiction between the two groups (P=0.531). These findings suggest that game-based training has a substantial impact on performance, self-efficacy, and motivation in adolescent non-athlete girls and may serve as an effective training strategy. The type of training method, whether game-based or traditional, does not appear to be a determining factor in the development or prevention of exercise addiction tendencies.

## 1. Introduction

The primary objective in sports coaching is to train and prepare athletes through effective and efficient methods. This goal is particularly significant in sports like futsal, where the unique conditions and characteristics of the game necessitate specialized training approaches. Futsal, a fast-paced indoor variant of football, demands players who possess a blend of technical skills, tactical awareness, and physical fitness to excel (1, 2). Given the inherent nature of futsal, with its high intensity, intermittent play, and emphasis on technical and tactical skills, exploring diverse training approaches to optimize player development is imperative (1). Performance in futsal is multifaceted, encompassing technical skills, tactical understanding, physical fitness, and psychological preparedness. Self-efficacy, defined as an individual’s belief in their ability to perform tasks and achieve goals, has been shown to have a significant relationship with futsal performance. Players with higher self-efficacy tend to demonstrate greater persistence, effort, and resilience when faced with challenges (3). Similarly, motivation plays a crucial role in futsal performance, influencing players’ commitment to training, desire to improve, and overall enjoyment of the sport (4). Conversely, exercise addiction, although less frequently studied, can potentially impact performance and lead to overtraining, burnout, or neglect of other important aspects of life (5). To develop these vital aspects of player performance, coaches employ various training methods and pedagogical strategies. Traditionally, sports training has relied heavily on structured, coach-centered approaches that emphasize repetitive drills and specific skill practice (6). This method, often referred to as the traditional training approach, focuses on breaking down complex skills into smaller, manageable components and practicing them in isolation before integrating them into game-like situations (6).In recent years, alternative strategies such as game-based learning have gained prominence in sports coaching (7). Teaching Games for Understanding (TGfU), proposed by Bunker and Thorpe in 1982, is an alternative approach that emphasizes the game itself, posing tactical and strategic problems in a modified game environment (8, 9). The TGfU model inverts traditional learning sequences: learners first develop a good understanding of the game, recognize its unique problems, and then practice the necessary skills to improve game performance. This approach integrates tactical and skill perspectives, emphasizing how tactics and skills should be taught according to the learner’s needs (10).

A comparison of traditional and game-based teaching methods reveals distinct differences in their approach and potential outcomes. While traditional methods may be more effective in developing specific technical skills through repetition, game-based approaches offer more holistic development by integrating physical, technical, tactical, and cognitive elements (11). Game-based learning is rooted in situated learning theory, which posits that learning is most effective when it occurs within authentic contexts. In futsal, this translates into training sessions that closely mimic real-game situations, allowing players to develop skills and make decisions under match-like conditions. This approach enhances technical and tactical proficiency while promoting the development of cognitive skills such as anticipation, decision-making, and problem-solving (12, 13). Research has shown that game-based approaches can lead to improvements in physical fitness, technical skills, and tactical awareness in futsal players (14). Additionally, these methods have been associated with increased motivation and enjoyment among participants, potentially leading to long-term participation in the sport (12). When comparing traditional and game-based teaching methods in relation to performance, self-efficacy, motivation, and exercise addiction, the literature presents mixed findings. Some studies suggest that game-based approaches may be more effective in developing decision-making skills and enhancing intrinsic motivation, while traditional methods have demonstrated advantages in acquiring specific technical skills and physical fitness (15). Despite the growing body of research supporting game-based training methods, there remains a gap in the literature regarding their effects on adolescent non-athlete girls in the context of futsal. Moreover, many existing studies have focused on male participants or athletes with prior sports experience (16, 17). Addressing this research gap could provide a broader understanding of game-based learning in non-athlete girls, ultimately allowing for a more comprehensive examination of the effects on motivation and exercise addiction alongside performance and self-efficacy.

Therefore, the aim of this study was to investigate the effects of 12 weeks of futsal-based game training on performance, self-efficacy, motivation, and exercise addiction in adolescent non-athlete girls.

## 2. Methodology

### 2.1 Data Collecting

This study employed a quasi-experimental design with a pre-test and post-test. The population consisted of all 13-17-year-old girls in Piranshar City. From this population, 30 individuals aged 13-17 (mean age 15) were randomly selected and assigned to two groups of 15: traditional training and game-based training. Inclusion criteria were physical and mental health, written parental consent, and no prior futsal training. Exclusion criteria included inability to perform skills, unwillingness to continue, and injury. The study was conducted in three phases: pre-test, intervention, and post-test.

In the pre-test phase, participants were familiarized with the exercises and research procedures, discussed training approaches separately, and the study’s objectives and dependent variables (performance (18), self-efficacy (19), motivation (20), and exercise addiction (21) were assessed. In the intervention phase, the research intervention was implemented using traditional and game-based training protocols for 12 sessions (1 hour each) separately for each group. The game-based training program was adapted from the study by Pizarro et al (2014), focusing on the principles of game modification, tactical awareness, and game simplification (22).

In the post-test phase, after the intervention, the dependent variables (performance (18), self-efficacy (19), motivation (20), and exercise addiction (21)) were reassessed. Performance included the assessment of futsal passing, shooting, and dribbling skills. In this study, all ethical principles of research were adhered to, including informed consent, confidentiality of information, and ensuring the health and safety of the participants. Table 1 shows the training protocol for the two groups.

**Table 1.** Training protocols by traditional training group and game practice training.

| session | Traditional Methods | Game Based training |
| --- | --- | --- |
| The first session | Warming up touching the ball, pass training, control and pass, pass to the wall, one-on-one pass, cooling down | pass training and its execution in 1 conditions close to the 2 x3game pass training and its 2 implementation in real game conditions |
| The second session | warm-up, touching the ball, one-on-one, pass under pressure of three people, one-hit and fast with defense, cool down | Training pass with pressure and 1 its implementation in conditions close to the game<br>Teaching pass with pressure and 2 its implementation in 1-2-1 game conditions |
| The third session | Warming up, reviewing the second session, shooting training, shooting against the wall with the dominant and non-dominant foot, shooting at the goal, cooling down | Pressure on attack, play on real pitch |
| The fourth session | Warming up, passing, shooting at the goal and combining passing and shooting, cooling down | Possession 0-4 |
| The fifth session | Warming up, passing, shooting, dribbling training on obstacles, dribbling with an assistant, cooling down | Progression sequence, dribble 1x1, 0-4 and 1-3 |
| The sixth session | Warming up, passing with pressure, shooting, dribbling under pressure, dribbling and shooting, cooling down | attack and defense |
| The seventh session | Warming up, dribble pass and shoot, pass and shoot, combine pass and shoot, cool down | Transition from attack to defense |
| The eighth session | Warming up, pass, shoot, dribble, combine pass, dribble and shoot, cool down | shoot |
| The ninth session | Warm-up, three-on-one passing to keep possession, four-on-two passing to keep the ball, shot with defensive wall, cool down | counterattack, retreat (delayed attack) |
| tenth session | Warming up, dribbling one against one and then shooting at the goal, passing and shooting three against two, cooled down | Attack tactics, 2x2 rotation, 1-2-1 system attack tactics |
| The eleventh session | Warming up, passing to the goal, shooting to the goal, dribbling between obstacles at maximum speed, cooling down | regional defense tactic 1-2-1 |
| The twelfth session | Warming up, playing with the aim of correct execution, passing, shooting, dribbling skills, cooling down | Combination of defense and attack tactics |

### 2.2 Futsal Skills Test

The Futsal Skills Test, designed by Marhaendro (2014), demonstrates high reliability and validity with coefficients of 0.89 and 0.95, respectively. The test is conducted on a marked 12-meter by 8-meter area of a futsal court. The setup includes:

- Two rebound zones (0.4 x 1 meters) on the longitudinal and transverse lines, marked with white, red, and yellow colors.
- Three passing zones (green, yellow, green), each measuring 1 meter square.
- Two dribbling zones for zigzag and pivot dribbling.
- Two 1-meter square shooting zones.
- A goal (3 meters wide, 2 meters high) divided into three 1 x 2 meter sections, with target zones marked in red near the goalposts.

Test Procedure:

1) The participant begins in the yellow passing zone, making 6 consecutive passes to rebound zone 1.
2) After completing the passes, they move to dribbling zone 1 for a pivot dribble, then return to the passing zone.
3) They then make 6 consecutive passes to rebound zone 2.
4) Next, they move to dribbling zone 2 for a zigzag dribble, returning to the starting zone.
5) Standing in the green zone near the transverse line, they make 6 consecutive passes to the rebound zone.
6) Finally, they move to the shooting zone and take two shots with their dominant foot and one with their non-dominant foot towards the target zone.
7) If all three shots hit the target zone, the test concludes. Otherwise, 4 additional shots are taken.

The test duration is recorded from start to finish. Penalties are applied for various errors:

- Hand-ball: 3 seconds
- Off-target shots: 2 seconds
- Hitting the goalpost: 1 second
- Hitting the middle of the goal: 0.5 seconds
- Shooting outside the designated area: 1 second
- Ball or shoe hitting a cone: 1 second each
- Pivot dribble outside the designated area: 1 second
- Passing to the red or white rebound zone area: 0.5 and 1 second, respectively
- Passing outside the rebound zone: 1 second
- Receiving the ball outside the designated area: 1 second

The total time is calculated by adding the performance time and penalty time. Each participant performs the test twice, with the best time recorded as their score (18).

### 2.3. Self-Efficacy

To assess participants’ self-efficacy, we employed the Physical Self-Perception Profile (PSPP) developed by Marsh and Richards (1982). This 22-item questionnaire is designed to measure individuals’ perceived physical self-efficacy in sports and physical activities. The PSPP comprises two subscales: The Perceived Physical Ability (PPA) subscale with 10 items and the Perceived Physical Condition (PPC) subscale with 12 items (19). Participants responded to each item on a 6-point Likert scale ranging from 1 (strongly disagree) to 6 (strongly agree). The PPA subscale assessed participants’ perceptions of their physical abilities, strength, and speed, while the PPC subscale measured their confidence in displaying physical skills in front of others. The overall Physical Self-Efficacy (PSE) score was calculated by summing all 22 items, with higher scores indicating greater physical self-efficacy. The validity and reliability of the PSPP have been well-established in previous research. Marsh and Richards (1982) reported high internal consistency for the overall PSPP scale (Cronbach’s α = 0.81) and for the PPA (α = 0.84) and PPC (α = 0.74) subscales. Test-retest reliability over a 4-week period was also satisfactory: PSE (r = 0.80), PPA (r = 0.85), and PPC (r = 0.69). The construct validity of the PSPP was supported by its correlations with related criteria, such as the Tennessee Self-Concept Scale and the Bem Sex Role Inventory (19). Additionally, the PSPP has demonstrated predictive validity in various sports and physical activity contexts (23, 24). For our study, we translated the PSPP into Persian using the back-translation method to ensure linguistic equivalence. The translated version was then pilot-tested with a small sample of adolescent girls. The reliability of this test in Iran was calculated using a test-retest method with a two-week interval in 43 students, resulting in a coefficient of 0.81 for the overall test. Its internal consistency was calculated with Cronbach’s alpha of 0.67 and a split-half reliability coefficient of approximately 0.72 (25).

### 2.4. Motivation

To assess participants’ motivation in sports contexts, we utilized the Sport Motivation Scale-II (SMS-II), developed by Pelletier and colleagues (2013). The SMS-II is an updated version of the original Sport Motivation Scale (20), designed to address previous limitations and better align with contemporary motivational theories, particularly Self-Determination Theory (SDT) (26). The SMS-II consists of 18 items divided into six subscales: intrinsic motivation, mixed regulation, self-regulated motivation, introjected regulation, external regulation, and amotivation, with each subscale containing three items. Participants responded to each item on a 7-point Likert scale ranging from 1 (not at all true) to 7 (completely true). Scoring for the SMS-II involved calculating the mean total score for each subscale by averaging the scores of the three items within that subscale (27). In initial validity assessments, the SMS-II demonstrated good internal consistency, with Cronbach’s alpha for the subscales ranging from 0.70 to 0.88 (27). Test-retest reliability was also reported as satisfactory (27). In terms of construct validity, the SMS-II showed expected correlations with related constructs. Intrinsic motivation was positively correlated with positive outcomes such as enjoyment and effort, while amotivation was negatively correlated with these outcomes and positively correlated with negative outcomes such as burnout. Additionally, different types of motivation exhibited correlations consistent with Self-Determination Theory (27).

### 2.5. Exercise Addiction

To assess exercise addiction among participants, we employed the Exercise Addiction Inventory (EAI) designed by Terry et al. (2004) (21). This questionnaire consists of six items, each corresponding to one of the six components of behavioral addiction: salience, mood modification, tolerance, withdrawal symptoms, conflict, and relapse (21). Participants responded to each item on a 5-point Likert scale, ranging from 1 (strongly disagree) to 5 (strongly agree). Scoring involved summing the scores of all six items, resulting in a total score ranging from 6 to 30. According to Terry et al. (2004), a score of 24 or higher indicates a risk of exercise addiction, a score of 13-23 suggests symptoms of exercise addiction, and a score of 0-12 indicates no exercise addiction. The EAI demonstrated good reliability in its initial validation study, with an internal consistency (Cronbach’s alpha) reported at 0.84, indicating strong reliability. Test-retest reliability was also satisfactory, with a correlation of 0.85 over a two-week period (28). In terms of construct validity, the EAI showed expected correlations with related constructs, positively correlating with the frequency and duration of exercise, as well as with measures of dependence and commitment to exercise (28). Furthermore, the EAI demonstrated concurrent validity, showing significant correlations with other established measures of exercise dependence, such as the Exercise Dependence Scale (29).

### 2.6 Statistical Analysis

Descriptive statistics were used to categorize the data. The Shapiro-Wilk test was used to examine the normality of the data distribution, and the analysis of covariance (ANCOVA) was used to examine the comparative effects of training. These steps were performed using SPSS version 26 at a significance level of 0.05.

## 3. Results

Table 2 shows the descriptive statistics of research variables separately for each group.

**Table 2.** Descriptive research findings.

| group variable | Traditional education |  | Game-oriented education |  |
| --- | --- | --- | --- | --- |
|  | pre-test | After the test | pre-test | After the test |
| **performance (total score)** | 53.39 ± 164.15 | 60.53 ± 156.11 | 27.64 ± 167.17 | 80.65 ± 141.9 |
| **self-efficacy (total score)** | 33.49 ± 11.3 | 13.2 ± 93.21 | 10.2 ± 20.78 | 1.17 ± 53.40 |
| **sports motivation (total score)** | 70.11 ± 53.58 | 14.82 ± 18.67 | 47.47 ± 72.14 | 60.51 ± 99.9 |
| **Sports addiction (total score)** | 67.55 ± 20.3 | 60.61 ± 23.2 | 20.2 ± 20.27 | 93.21 ± 22.2 |

To assess the normality of the pre-test data distribution, the Shapiro-Wilk test was employed. The results indicated that for all variables, the significance level was greater than 0.05, confirming the assumption of a normal distribution of the data (P > 0.05). Consequently, parametric tests were utilized to analyze the data in testing the research hypotheses. To examine the effect of the intervention on the research variables, analysis of covariance (ANCOVA) was selected. Prior to conducting this analysis, the assumptions of homogeneity of variance and equality of regression slopes were assessed and confirmed. Levene’s test indicated that the assumption of homogeneity of variances was satisfied for all research variables (P > 0.05). Therefore, after controlling for the effect of pre-test scores, post-test scores between the two groups were compared using ANCOVA, with results presented in Table 3.

**Table 3.** Analysis of covariance test results.

| Variable | Statistical effect | sum of squares | df | Mean square | F | Sig | Effect size |
| --- | --- | --- | --- | --- | --- | --- | --- |
| **performance** | covert | 2125.63 | 1 | 2125.68 | 55.05 | 0.0001 | 0.67 |
|  | between groups | 1963.33 | 1 | 1963.33 | 50.85 | 0.0001 | 0.65 |
| **self-efficacy** | covert | 63.93 | 1 | 63.93 | 52.73 | 0.0001 | 0.66 |
|  | between groups | 124.37 | 1 | 124.37 | 102.59 | 0.0001 | 0.79 |
| **Sports motivation** | covert | 2333.09 | 1 | 2333.09 | 36.00 | 0.0001 | 0.57 |
|  | between groups | 1811.61 | 1 | 1811.61 | 95.27 | 0.0001 | 0.50 |
| **Sports addiction** | covert | 23.51 | 1 | 23.51 | 44.4 | 0.044 | 0.14 |
|  | between groups | 2.04 | 1 | 2.04 | 0.39 | 0.537 | 0.01 |

According to table 3. The results of covariance analysis showed that there was a significant difference in performance (P=0.0001), self-efficacy (P=0.0001) and sports motivation (P=0.0001) between the game-based training group and the traditional training group. There was no difference in exercise addiction between the two groups (P=0.531).

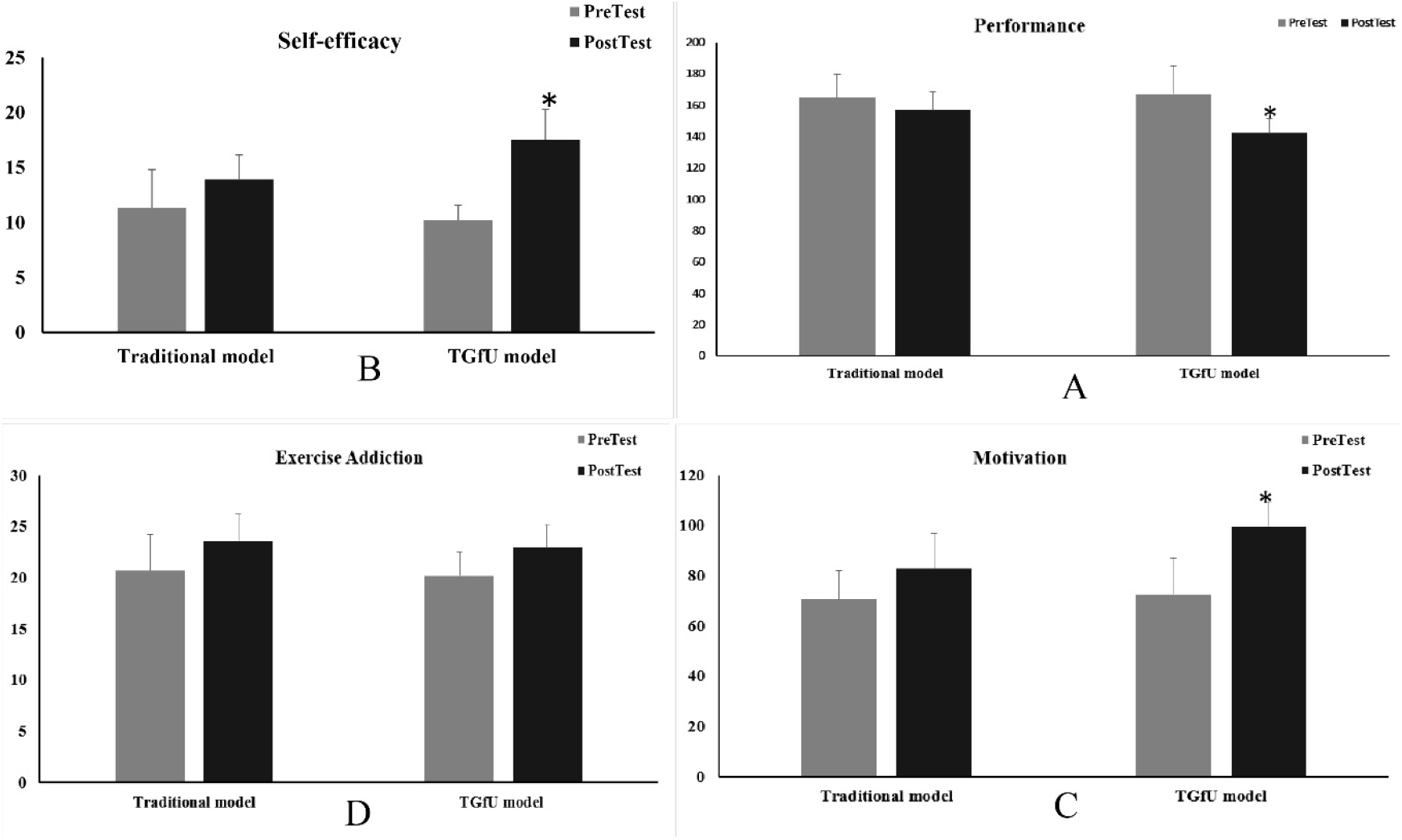

test conditions in terms of performance (A) self-efficacy (B) sports motivation (C) sports addiction (D) * significant difference between the traditional education group and the game-based education group at the level of P<0.05)

## 4. Discussion

The present study aimed to investigate the effects of a 12-session futsal training program on the performance, self-efficacy, motivation, and exercise addiction of non-athletic adolescent girls. The implementation of game-based training strategies in sports, particularly in futsal, has garnered significant attention in recent years due to its potential to enhance various aspects of athletic development. Futsal, a fast-paced variant of indoor soccer, is recognized for its ability to improve technical skills, tactical awareness, and physical fitness across all age groups (15). The unique characteristics of the game—such as a smaller playing area, frequent ball contacts, and the need for quick decision-making—make it an ideal platform for developing fundamental soccer skills and overall athletic abilities (30). Game-based training strategies in futsal are particularly relevant given the multifaceted nature of athletic development. Improved performance in sports is not solely dependent on physical attributes but is also influenced by psychological factors such as self-efficacy and motivation (31). Self-efficacy, defined as an individual’s belief in their ability to perform specific tasks, plays a significant role in sports performance and sustained participation (32). Similarly, both intrinsic and extrinsic motivation are key determinants of sports engagement and long-term adherence to physical activity (6). Additionally, the concept of exercise addiction, while less explored in youth sports literature, warrants attention due to its potential impact on overall well-being and the development of a balanced lifestyle (33). The results of the present study indicated that game-based training significantly impacted performance, self-efficacy, and sports motivation. Previous studies have similarly demonstrated that game-based training can effectively enhance performance (34-36), aligning with our findings. However, some studies have reported inconsistent results (37-40). In these studies, training based on game-based teaching styles had a direct impact on motor skill performance and certain psychological characteristics, while other studies found no significant differences compared to traditional methods. In team sports like futsal, which involve open skills, players must constantly coordinate their actions with teammates and opponents to achieve optimal performance outcomes. This requires quick decision-making under the constraints of the game environment. From an ecological dynamics perspective, tactical behavior is an active and ongoing process of searching for and exploring information on the playing field to guide performance (37-40). To enhance players’ tactical behavior, it is crucial to design training exercises that expose them to game-like conditions. This forms the basis of game-based training, which replicates the perceptual-motor demands of real competition. The game-based learning approach emphasizes manipulating task constraints to simplify game situations, support limited information processing, and guide players toward primary goals (22). This concept aligns with Newell’s constraint-based approach (41), which posits that manipulating performance constraints during training can effectively prepare individuals for real competitive conditions. In our study, the game-based training program likely improved performance by manipulating constraints similar to those encountered during competition. However, it is important to note that differences in findings across studies may be attributed to factors such as the limited duration of training, the specific sport, or the skill level of the participants (37-40). These factors should be carefully considered when interpreting the effectiveness of game-based training in various sports contexts.

The findings of the present study regarding the impact of game-based training on physical self-efficacy align with those of Wilang et al. (35), who demonstrated that children exhibited significantly higher self-efficacy when participating in game-based programs compared to traditional physical education. Additionally, a study examining teachers’ self-efficacy in educational play found that physical education teachers reported higher self-efficacy than preschool teachers. This suggests that game-based learning can specifically enhance self-efficacy in physical education contexts (42).

In the context of our study, the significant improvement in physical self-efficacy among non-athletic adolescent girls participating in futsal-based training can be attributed to several factors. The nature of futsal, characterized by fast-paced gameplay and frequent ball contacts, likely provides numerous opportunities for success and skill development. The progressive structure of the 12-session program may have allowed participants to experience gradual improvements, thereby increasing their confidence over time. Furthermore, the team-oriented nature of futsal may have fostered a supportive environment, enhancing verbal persuasion and vicarious learning experiences (43).

It is important to note that our study focused on non-athletic adolescent girls, a population that may have lower baseline levels of physical self-efficacy compared to experienced athletes. This lower baseline could create more room for improvement, potentially explaining why our results align more closely with studies demonstrating positive effects rather than those reporting no significant changes. The positive impact of game-based training on self-efficacy can be understood through several mechanisms rooted in Bandura’s self-efficacy theory (43). Firstly, game-based training offers numerous opportunities for mastery experiences, which are considered the most influential source of self-efficacy. As participants successfully execute skills and make decisions in realistic contexts, their belief in their abilities is strengthened. Secondly, vicarious experiences play a role, as players observe their peers’ successful performances during training, further enhancing their self-efficacy. Verbal persuasion from coaches and teammates, in the form of encouragement and positive feedback, also contributes to this enhancement. Lastly, the physiological and affective states associated with the excitement of game situations may positively influence self-efficacy beliefs (43). However, it is essential to acknowledge that not all studies have reported positive effects of game-based training on self-efficacy (40). These contradictory findings highlight the complexity of self-efficacy development and suggest that various factors may influence the effectiveness of game-based training interventions. Several explanations for the lack of effectiveness in some studies have been proposed. One study suggested that ceiling effects may occur when participants already possess high baseline levels of self-efficacy, thereby limiting the potential for improvement (40). Additionally, the specific sport and the design of game-based training programs may significantly influence their effectiveness in enhancing self-efficacy (40).

Here’s a revised version of your discussion on the effectiveness of game-based training on sports motivation and exercise addiction, ensuring clarity, coherence, and adherence to scientific writing standards:

The results of our study regarding the effectiveness of game-based training on sports motivation align with several studies that have demonstrated a positive impact of such training. For instance, Allison and Thorpe conducted a study comparing the effects of game-based and skill-based approaches on motivation in physical education classes, finding that students in the game-based group exhibited higher levels of motivation and intrinsic enjoyment compared to those in the skill-based group (44). Similarly, Gray et al. examined the effects of game-based training on the motivation of young soccer players, revealing that participants in the game-based training group reported higher levels of motivation and perceived competence than those in the traditional training group (45). Gomes et al. conducted a review study that highlighted the positive effects of game-based learning and gamification strategies in physical education, noting their impacts on student engagement, academic achievement, and motivation in sports contexts.

However, not all studies have reported positive effects of game-based training on sports motivation. For example, Caglar and Asci investigated the effects of a game-based physical education program on students’ motivation and found no significant differences between the experimental and control groups (46). Interpreting our results in light of these previous studies, the significant improvement in sports motivation observed in our futsal-based training program for non-athletic adolescent girls is more consistent with the findings of Allison (44) and Gray et al. (45). This consistency suggests that game-based training, particularly in team sports like futsal, can effectively enhance motivation in young athletes.

The positive results may be attributed to several factors inherent in game-based training approaches, such as increased enjoyment, perceived autonomy, and opportunities for social interaction (47, 48). The effectiveness of game-based training in enhancing sports motivation can be explained through mechanisms rooted in Self-Determination Theory (SDT). Game-based training often fulfills the three basic psychological needs outlined in SDT: autonomy, competence, and relatedness. The dynamic nature of game-based activities allows participants to make decisions and solve problems independently, fostering a sense of autonomy. The progressive challenges and immediate feedback provided in game situations contribute to feelings of competence as participants experience success and progress. Additionally, the team-oriented nature of futsal promotes social interaction and cooperation, fulfilling the need for relatedness (36, 49).

The lack of effectiveness reported in some studies may be attributed to various factors. The duration of the intervention plays a crucial role in inducing motivational changes; short-term interventions may not provide sufficient time to produce significant changes in motivation. Moreover, the initial motivation levels of participants can influence their potential for improvement. Those with already high motivation may show limited progress due to ceiling effects. The specific sport and the design of game-based training programs may also affect their effectiveness in increasing motivation (50).

Regarding the lack of significant differences between the traditional training group and the game-based training group in terms of exercise addiction, our findings align with the research of Lichtenstein et al. (51). Their study comparing the effects of different training methods on exercise addiction in recreational runners found no significant differences in exercise addiction between groups of football players and bodybuilders who exercised individually. Marquez et al. (52) showed that the prevalence of exercise addiction symptoms varies across sports and competition levels, with elite athletes in sports such as bodybuilding, weightlifting, and rowing exhibiting higher levels of exercise addiction compared to other sports. The authors suggest that the nature of the sport and the competitive environment may influence the risk of developing addictive exercise behaviors. However, not all studies have reported similar results. Sabo et al. (53) conducted a brief review analysis that reported a significant prevalence of exercise addiction risk in elite athletes compared to those who exercise for leisure. Some research has indicated differences between game-based and traditional training approaches in factors that may influence exercise addiction. Fortier et al. found that participants in a game-based training program exhibited higher levels of intrinsic motivation and more self-determined forms of behavioral regulation compared to those in a traditional training program, which could potentially influence the risk of developing exercise addiction (54). Additionally, McNamara and McCabe (55) examined the relationship between training approaches and exercise dependence in competitive athletes. Their results indicated that athletes participating in more structured and traditional training programs were at a higher risk of developing exercise dependence compared to those engaged in more varied and game-based training approaches (55). Interpreting our results in light of these previous studies, the lack of significant difference in exercise addiction between the game-based and traditional training approaches in our study of non-athletic adolescent girls aligns with the findings of Lichtenstein et al. (51). This consistency suggests that the type of training approach may not be the primary factor in developing or preventing exercise addiction, at least in the context of a short-term intervention with non-athletic individuals. The absence of differences in exercise addiction between the two training approaches can be explained through various mechanisms. Firstly, exercise addiction is a complex phenomenon influenced by multiple factors, including personality traits, psychological needs, and environmental factors, which may not be significantly altered by a particular training approach over a short period. Secondly, both game-based and traditional training approaches can provide opportunities for skill mastery, social interaction, and physical activity—key elements that contribute to the beneficial aspects of sports participation (56, 57). Finally, the duration of our intervention (12 sessions) may not have been sufficient to induce measurable changes in addictive tendencies related to sports participation.

The inconsistent results reported in some studies can be attributed to various factors. Different training approaches may influence motivational and behavioral regulation factors, which can indirectly affect the risk of exercise addiction over time (58). Additionally, the study population—whether competitive athletes or non-athletes—may play a significant role in the potential for developing exercise addiction. Furthermore, the duration and intensity of the intervention, as well as the specific measures used to assess exercise addiction, may also influence the observed results (58). The absence of a significant difference in exercise addiction between game-based and traditional training approaches in non-athletic adolescent girls suggests that the type of training may not be a critical factor in the short-term development or prevention of exercise addiction. This study examined the effects of a 12-session futsal-based training program on performance, self-efficacy, motivation, and exercise addiction in non-athletic adolescent girls. While this research yielded interesting results, particularly in motivation and self-efficacy, it also faced several limitations that may have influenced its effectiveness or lack thereof in specific variables, especially exercise addiction.

The present study was subject to several limitations. Key limitations included the characteristics of the sample population, the relatively short duration of the intervention, and the absence of a non-intervention control group. Focusing on non-athletic adolescent girls limits the generalizability of the findings, while the 12-session program may not have been sufficient to observe changes in complex phenomena such as exercise addiction. The lack of a control group makes it challenging to attribute changes solely to the training program, as individual differences among participants and external factors could influence the results independently of the intervention. Overall, future research should investigate longer intervention durations, different types of interventions, and additional factors such as feedback to gain a broader understanding of this training strategy.

## 5. Conclusion

The findings indicate that game-based training leads to significant improvements in performance, self-efficacy, and sports motivation compared to traditional training methods; however, no difference was observed in exercise addiction. Based on the results, the engaging and dynamic nature of game-based training appears to be more effective in motivating participants and building confidence in their athletic abilities. Nevertheless, the type of training method—whether game-based or traditional—may not be a determining factor in developing or preventing tendencies toward exercise addiction, at least within the scope and duration of this study.

